# Synergism and phytotoxicity: the effects of tank-mix additives on the biological efficacy of Cu^2+^ against *Venturia inaequalis* and *Podosphaera leucotricha*

**DOI:** 10.1101/2022.11.08.515630

**Authors:** Christine Schmitz, Eike Luedeling, Shyam Pariyar

**Author notes:** Corresponding author: Shyam Pariyar.

## Abstract

The wetting behaviour of the spray and biological efficacy of Cu^2+^ active ingredients in agrochemical formulations may be enhanced by tank-mix additives. We investigated how three BREAK-THRU^®^ additives (BT301: biodegradable, BT133 and BT420: bio-based and biodegradable) as tank-mix with commercial copper preparations influence the spray distribution, leaf uptake and biological efficacy of copper additive mixtures against apple scab and apple powdery mildew under controlled conditions. We quantified the synergetic effects of these additives in foliar applications. In addition, we determined the phytotoxic potential and evaluated impacts on photosynthetic activity, non-photochemical quenching and ROS activity. The additives BT301 and BT420 strongly reduced surface tension and contact angle of copper treatments. The fluorescence observations revealed that BT301 achieved better spreading of copper formulation with more complete coverage of the leaf surface than BT420 and BT133, whereas “coffee-ring” spreading was observed with BT133. The additive BT301 showed an increase in relative fluorescence area, indicating higher ROS production as a signal of intra-cellular tissue activity. The photochemical efficiency of photosystem II (Fv/Fm) was not negatively influenced by copper or additive treatment. Thus, we observed no phytotoxic effects of copper-additive mixtures on apple leaves at treatment doses of 4 g Cu^2+^ L^-1^. All copper treatments reduced apple scab infestations significantly, by 53-76%. Interestingly, addition of BT301 to copper preparations showed the strongest biological efficacy (83% reduction) against *V. inaequalis*, whereas addition of BT420 showed the strongest effect against *P. leucotricha* (89% infection reduction). The synergetic effects of additives on the biological efficacy without phytotoxic effects on plants may have potential for reducing copper loads in horticultural production systems.

## 1. Introduction

The apple scab pathogen *Venturia inaequalis* and the apple powdery mildew pathogen *Podosphaera leucotricha* are two important fungal pathogens in apple cultivation regions all over the world (Holb et al., 2017). *V. inaequalis* has an exceptional life strategy that does not involve host cell penetration. Instead, this fungus develops in the subcuticular space until it sporulates (Bowen et al., 2011). *P. leucotricha* penetrates the leaf epidermal cells, forming haustoria that extract nutrients from the host plant (Marine et al., 2010).

Foliar application of copper fungicides is a widespread practice in agriculture. Copper is widely used both in conventional and organic farming systems worldwide (Ostandie et al., 2021). High copper accumulation in soil and aquatic environment is harmful to humans and other organisms and increases ecological risks (Rehman et al., 2019). Thus, copper application has become an ecological and political concern in many places and its application in organic production has already been restricted in some European countries (Holb and Kunz, 2016). Since there is no complete alternative or economic replacement for copper use in organic farming, reducing copper loads in horticultural production systems is seen as a potential strategy to avoid copper accumulation on farmland (Holb and Kunz, 2016; Kuehne et al., 2017; Tamm et al., 2022). Copper acts as a broad-spectrum biocide and is used to control several pathogens, including *V. inaequalis*, in Integrated Pest Management as well as in organic production systems (Lamichhane et al., 2018; Kuehne et al., 2017). As an essential nutrient element, copper has important functions within plants, e.g. for photosynthesis. However, copper is also a heavy metal that can be phytotoxic at high concentrations, and frequent applications can lead to its accumulation in the soil (Rusjan, 2012). Copper application to apple leaf surfaces can help prevent *V. inaequalis* infection by suppressing spore germination and appressorium formation of the apple scab disease (Montag et al., 2006). A post-infection spray of copper hydroxide can also be effective, but effectiveness depends on how much time has passed since the initial infection. After 24 h, copper has been shown to kill existing stromata and prevent formation of new stromata by *V. inaequalis*, but this effect was no longer observed after 48 h (Montag et al., 2006). Copper hydroxide particles need a water film in contact with the fungal structures to be effective (Montag et al., 2006). However, copper accumulations on plant leaves can be phytotoxic (Lalancette & McFarland, 2007).

Tank-mix additives interact with the agrochemicals, modifying the chemicals’ mode of action or the physical properties of the mixture (Hazen, 2000). Added to pesticides or foliar fertilisers, tank-mix additives may improve effectiveness (Melo et al., 2015). In addition, some additives or tank-mix formulations could have phytotoxic effects on crop plants (Burchill et al., 1979; Butt et al., 1973; Räsch et al., 2018). Several tank-mix additives of the BREAK-THRU^®^ series, such as BREAK-THRU^®^ S 233 and S 240, are known to reduce the surface tension and contact angles of different agrochemicals (Basi et al., 2014; Melo et al., 2015). However, there is little information to date on the synergistic effects of these additives on copper-pathogen interactions. The aim of the present study was to investigate three BREAK-THRU^®^ tank-mix additives (BT301: biodegradable, BT133 and BT420: bio-based and biodegradable) to evaluate whether i) they have synergistic effects on copper treatments against *V. inaequalis*, the causal agent of apple scab, and *P. leucotricha*, which causes apple powdery mildew, ii) they are phytotoxic after foliar application and iii) to elucidate the physical, chemical and physiological mechanisms behind the above interactions.

## 2. Materials and Methods

### 2.1 Plant material and growth conditions

Apple seedlings (*Malus domestica*, cultivar *Bittenfelder*) were grown in a protected environment under controlled growth conditions (19.44 ± 0.02 °C, 73.92 ± 0.10% RH) in plastic containers (volume 1 liter, Teku^®^ VCG 14, Pöppelmann, Germany). The containers were filled with a peat mixture substrate consisting of peat, sand and perlite at a mixing ratio 1:1.2:0.3. The substrate had a bulk density of 0.57 g cm^-3^ and contained 318 g of water at 60% water holding capacity. The plants were irrigated automatically 1-2 times per day with a drip irrigation system, depending on water demand, maintaining soil moisture at 54-64% of water holding capacity. Plants were supplied with full nutrition with an osmocote slow release NPK (Mg) fertiliser with micro-nutrients (“Osmocote Exact”; Everris, The Netherlands). Light (145 ± 14 μmol m^-2^ s^-1^ photoactive radiation, PAR at canopy level) was provided for 10 hours per day (8 a.m. to 6 p.m.). Five plants for each treatment were placed as randomly designed blocks under the same growth conditions. Each treatment block (five plants) were re-positioned within the experimental setting twice per week to maintain equal exposure to growth conditions during the experiments.

### 2.2 Treatments

We used three BREAK-THRU^®^ tank-mix additives: i) BT133, a bio-based (produced with renewable resources, containing 95% bio-based carbon) and biodegradable sticker penetrant based on polyglycerol esters and fatty acid esters, ii) BT301, a biodegradable super spreader based on an organo-modified polyether trisiloxane, and iii) BT420, a bio-based and biodegradable spreader based on glycolipids (Evonik Operations GmbH, Germany). The dosage was calculated based on the recommended rates for orchards (Table 1). As sources of copper fungicides, we used three copper hydroxide (Cu(OH)_2_) formulations: Funguran^®^ progress (wettable powder), Cuprozin^®^ progress (suspension) (Certis Europe B.V., Germany) and TEGO^®^ XP 11052 (suspension) (Evonik Operations GmbH, Germany). Copper treatments were applied either alone or in a mixture with one tank-mix additive. As control treatment, we used demineralised water.

**Table 1:** List of selected treatment products and dosages (diluted with demineralised water). Additives: BREAK-THRU^®^ SP 133 (BT133), BREAK-THRU^®^ S 301 (BT301) and BREAK-THRU^®^ SF 420 (BT420). Copper treatments: Funguran^®^ progress (Fung), Cuprozin^®^ progress (Cup) and TEGO^®^ XP 11052 (TEGO_Cu). Units in concentration are, v/v = volume/volume or g L^-1^.

| Treatment | Concentration | Ca<sup>2+</sup> or Cu<sup>2+</sup> [mg L<sup>-1</sup>] |
| --- | --- | --- |
| BT133 | 0.015 % v/v | - |
| BT301 | 0.008 % v/v | - |
| BT420 | 0.025 % v/v | - |
| Fung |  |  |
| (Copper hydroxide fungicide, Wettable powder formulation) | 3 g L<sup>-1</sup> | 1050 |
| Cup | 2 g L<sup>-1</sup> | 375 |
| (Copper hydroxide fungicide, suspension formulation) | 4 g L<sup>-1</sup> | 750 |
|  | 12 g L<sup>-1</sup> | 2251 |
| TEGO_Cu | 2 g L<sup>-1</sup> | 375 |
| (Copper hydroxide fungicide, suspension formulation) | 4 g L<sup>-1</sup> | 750 |
|  | 12 g L<sup>-1</sup> | 2251 |

### 2.3 Surface tension and contact angle

Contact angle and surface tension were measured with a drop shape analyser (DSA30; Krüss GmbH, Germany) at room conditions (23.14 ± 0.13 °C, 45.00 ± 0.34% RH). Contact angles of 10 droplets (volume, 5 μl) per treatment were measured using the sessile drop method. Contact angles were determined on a silicone surface (as smooth model surface), as well as on the adaxial (upper side, astomatous) and abaxial (lower side, stomatous) sides of apple leaves. Leaves were obtained from similar positions within the trees (16-20th leaf, counting from the bottom of the canopy). To ensure a level leaf surface, we attached the leaves to specimen slides using double-faced adhesive tape (tesa SE, Germany), taking care not to disturb the measurement surfaces. Since left and right contact angles of the droplets were symmetrical and showed no significant differences (P > 0.05, n = 10), they were merged together for the treatment group comparison. We measured the surface tension of 10 droplets per treatment using the pendant drop method.

### 2.4 Surface coverage of copper preparations and intra-cellular tissue activity

Shape and distribution of deposited residues of copper formulations with and without selected BREAK-THRU^®^ treatments were visually analysed using a multispectral fluorescence imaging system (Nuance CRI, Perkin-Elmer, United Kingdom). Six droplets (volume 5 μl) of each treatment were applied to the adaxial side of an apple leaf. After the droplets had dried out, a fluorescence marker (2’,7’-dichlorofluorescin diacetate; H2DCFDA) prepared in a 50 mM phosphate buffer solution was misted onto treated leaves using a fingertip sprayer. H2DCFDA serves as an indicator of reactive oxygen species (ROS) signalling intra-cellular tissue activity (Fichman et al., 2019). After the fluorescence marker had dried out, RGB pictures (520-540 nm) were taken at 20x magnification (i.e. picture area: 36 mm^2^). Treatment solutions were also sprayed on the abaxial side of apple leaves using a fingertip sprayer. Five days after treatment application, the plants were misted with H2DCFDA. After the fluorescence marker had dried out, RGB pictures (420-640 nm) were taken at 11x magnification (i.e. picture area: 1.24 cm^2^). The relative fluorescence area, an indicator of intra-cellular tissue activity, was determined using the software ImageJ Version 1.53c (Wayne Rasband, National Institutes of Health, USA) running on Java Version 1.8.0_261 (Oracle, USA).

### 2.5 Chlorophyll fluorescence and phytotoxic potential

To evaluate whether the additives alone or as tank mixtures with copper hydroxide formulations have a phytotoxic effect on apple leaves, we measured leaf chlorophyll fluorescence. Leaves from similar positions within the plant and of similar age (1-2 weeks) were selected for phytotoxicity trials. Five leaves of each plant were treated either on the adaxial or the abaxial side. We applied four single droplets (at 5 μl per droplet) of each tested substance to the leaf using a Hamilton Repeating Dispenser (Sigma-Aldrich, Germany). Each treatment had five replicates, in a randomized design over plants and leaf position, to avoid antagonistic effects of leaf age on photosynthetic activity. The chlorophyll fluorescence measurements on apple leaves were done by pulse-amplitude modulated (PAM) chlorophyll fluorometry combined with saturating pulse analysis of fluorescence quenching. This measurement provides quantitative information on the quantum yield of photosynthetic energy conversion and photochemical stress responses (Klughammer & Schreiber, 2008; Pariyar & Noga, 2018). Chlorophyll fluorescence images (640 × 400 pixel) were taken 7 days after treatment on the abaxial leaf side and 8 days after treatment on the adaxial leaf side. Leaves were dark-adapted for at least 20 minutes before the measurement. The leaves were carefully positioned and attached to the frame of the IMAGING-PAM under near infrared light (780 nm) approximately 100 mm away from the radial light source. The induction curves were recorded under laboratory conditions (i.e. at a temperature of approximately 25 °C and ambient CO2 concentration) using an imaging PAM fluorometer (ImagingPAM, Heinz-Walz GmbH, Effeltrich, Germany), equipped with 96 blue light-emitting diodes (peak at 470 nm) used for fluorescence excitation, actinic illumination and saturation pulses. Circular areas of interest (AOI) were manually chosen with the ImagingWin software at the solution application sites. Ground fluorescence (Fo) was recorded after leaf illumination by the blue light-emitting diodes (0.5 μmolm^−2^ s^−1^); the maximum fluorescence (Fm) was determined after a blue light saturation pulse of 1000μmol m^−2^ s^−1^. The yield of variable chlorophyll fluorescence (Fv) was calculated as Fm-Fo, while the maximum photochemical efficiency of the (PSII) was calculated as Fv/Fm. After the first saturation pulse, actinic light (225 μmol m^−2^ s^−1^) was switched on and saturation pulses (1000 μmol m^−2^ s^−1^) were applied at 30 s interval within a period of 175 s. We also quantified additional photochemical stress response parameters, such as maximum fluorescence yield (Fm’), quantum yield of regulated energy dissipation in PSII, Y(NPQ), quantum yield of non-regulated energy dissipation in PSII, Y(NO) and apparent rate of photosynthetic electron transport (ETR, μmol m^−2^ s^−1^).

### 2.6 Biological efficacy

Thirty apple seedlings of similar age and condition were used for a biological efficacy trial, with 4 leaves from each plant selected for treatments and efficacy rating. The plants were pre-treated either with demineralised water as control or with copper preparations with and without BREAK-THRU^®^ additives. Sprays were applied to five plants per treatment using a plant sprayer (MESTO GmbH, Germany). After the treatments had dried out, an inoculum (about 430,000 conidia ml^-1^) of apple scab (*V. inaequalis*) conidia was sprayed onto all plants (about 1.67 ml per plant) with a plant sprayer. The inoculated plants were incubated in a box at room temperature (23.73 ± 0.05 °C) and high humidity (99.60 ± 0.08 % RH) for 48 hours to allow the spores to germinate. After incubation, the plants were returned to the growth chamber, where they were maintained at a temperature of 20.01 ± 0.03 °C and 57.16 ± 0.12% RH during the day and 20.79 ± 0.04 °C with 68.22 ± 0.18% RH at night. No inoculation with *P. leucotricha* was done, since *P. leucotricha* infestation was expected to occur naturally in this growth environment. The apple seedlings were not pre-treated against *P. leucotricha* with other fungicides. So we were able to evaluate the synergistic effect of copper formulations on apple powdery mildew suppression. Thus, we evaluated plants for infestation by *V. inaequalis* and *P. leucotricha* 7 and 14 days after inoculation on the four selected leaves of each plant. Infestation of each leaf was estimated in percent of leaf area.

### 2.7 Statistical analysis

Statistical analysis was performed using the software R version 4.0.2 (R Core Team, 2020) and the packages tidyverse version 1.3.0 (Wickham et al., 2019) and agricolae version 1.3-3 (De Mendiburo, 2020). Results were stated as mean ± standard error (SEM). We evaluated the effects of the treatments using analysis of variance (ANOVA). The assumptions of an ANOVA were tested with Residual vs. Fitted and Normal Q-Q plots. LSD-tests (LSD 5%, least significant difference between two means at P = 0.05;) were conducted to screen for significant differences between groups. Paired t-tests were used for comparisons between individual treatment groups. For all statistical tests, the significance level was α = 0.05. Graphs were made with ggplot2 version 3.3.2 (Wickham, 2016).

## 3. Results

### 3.1 Surface tension and contact angles

The addition of tank-mix additives to demineralised water reduced the surface tension considerably by 2.7% with BT133, 46.0% with BT420 and 73.2%with BT301 (Figure 1a, Supporting Information Table S1). Adding BT420 significantly reduced the contact angles on apple leaf surfaces on both the adaxial (by 32.9%) and the abaxial (by 29.2%) side (Figure 1b, Supporting Information Table S2). For BT301, contact angle measurement on leaf surfaces was not possible, because the solution spread too fast. On the silicone surface, BT301 produced the strongest reduction in contact angles (Supporting Information Table S1).

**Figure 1:**
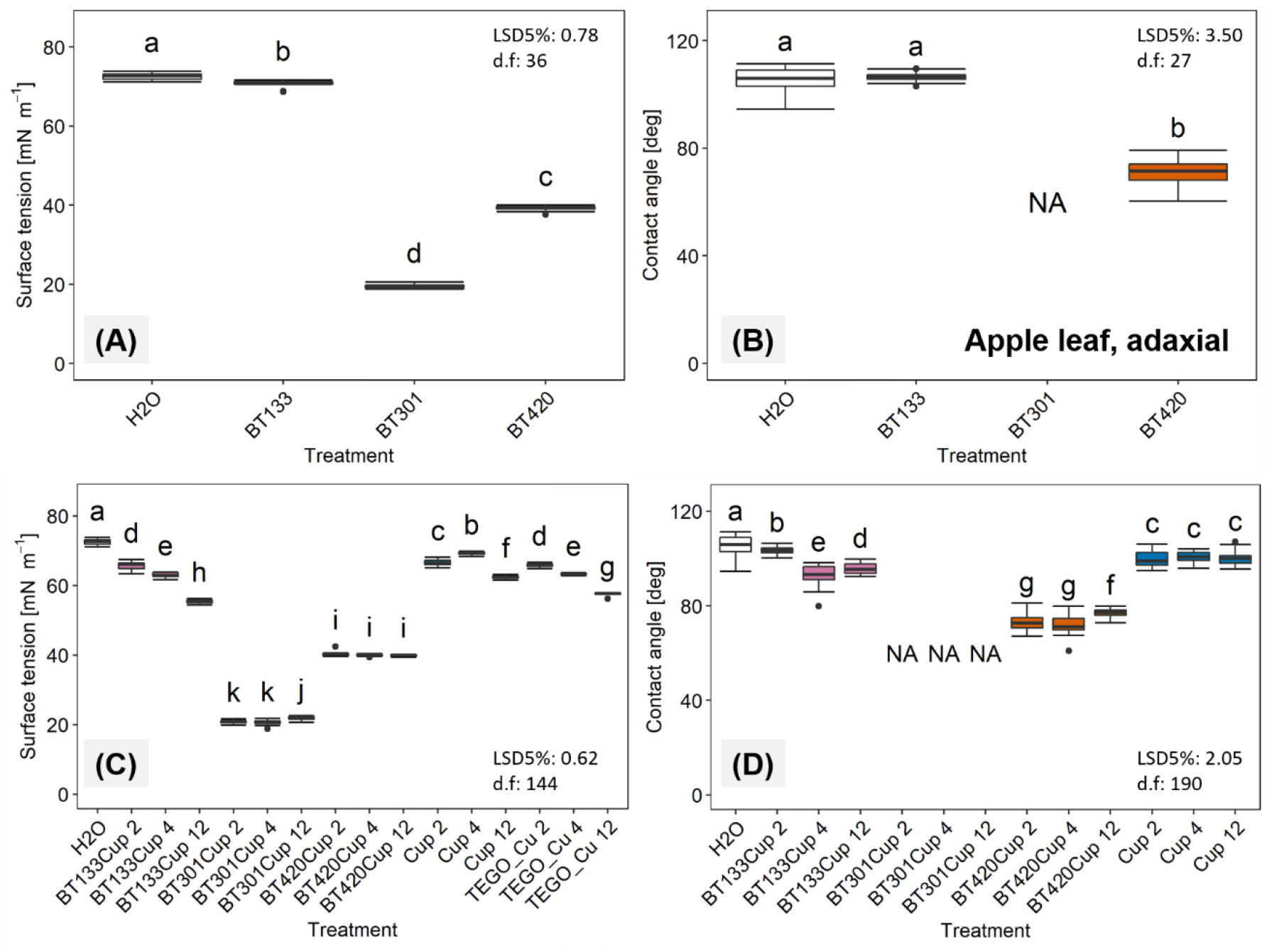
Surface tension [mN m^-1^] (a, c) and contact angles [°] on adaxial apple leaf surfaces (b, d) of different BREAK-THRU^®^ additives (a, b) and different doses (2, 4 and 12 g L^-1^) of copper preparations (c, d) with BREAK-THRU^®^ (BT133: 1.5 ml L^-1^, BT301: 0.75 ml L^-1^, BT420: 2.5 ml L^-1^) additives dissolved in water. Droplet size for contact angles was 5 μl, and left and right contact angles were merged. Boxplot shows median ± quartile (lower 25%, upper 75%), n = 20 for contact angles, and n = 10 for surface tension. Different letters denote significant differences between treatment groups (LSD, least significant difference between two means at P = 0.0; d.f., degrees of freedom associated with LSDs 5). NA = data are not available because the contact angle measurement of treatments containing BT301 was not possible on apple leaves due to fast spreading.

All copper formulations had significantly lower surface tension compared to demineralised water (Figure 1e, Supporting Information Table S3). We found the strongest effect for the addition of BT301, which reduced the surface tension of Fung and Cup by about 70% compared to demineralised water (Figure 1e, Supporting Information Table S3). The addition of BT420 reduced the surface tension by about 43% (Figure 1e). Contact angles of Cup and Fung were reduced by the additives (Figure 1f, Supporting Information Table S3).

### 3.2 Surface coverage of copper preparations

The visual analysis of surface coverage by different treatments showed that directly after application BT133 formed coffee-ring type droplets, which often merged due to leaf movement (Figure 2a). BT420 first covered a round area and dried out leaving small uncovered areas (Figure 2c). Droplets of Cup on apple leaves formed coffee-ring structures leaving potentially uncovered surface within the sprayed area (Figure 2b). The distribution of Cup on the leaf surface was improved by BT301 and BT420 (Figure 2d, 2f). The fluorescence observations suggested that BT301Cup achieved better spreading with more complete coverage of the leaf surface than BT420Cup, whereas BT420Cup produced a dense and even distribution of copper preparation within the droplets (Figure 2d, 2f). BT301 spread so quickly after application that a droplet shape could not be discerned. BT301 treatments produced this effect when applied alone or in combination with Cup (Figure 2e, 2f).

**Figure 2:**
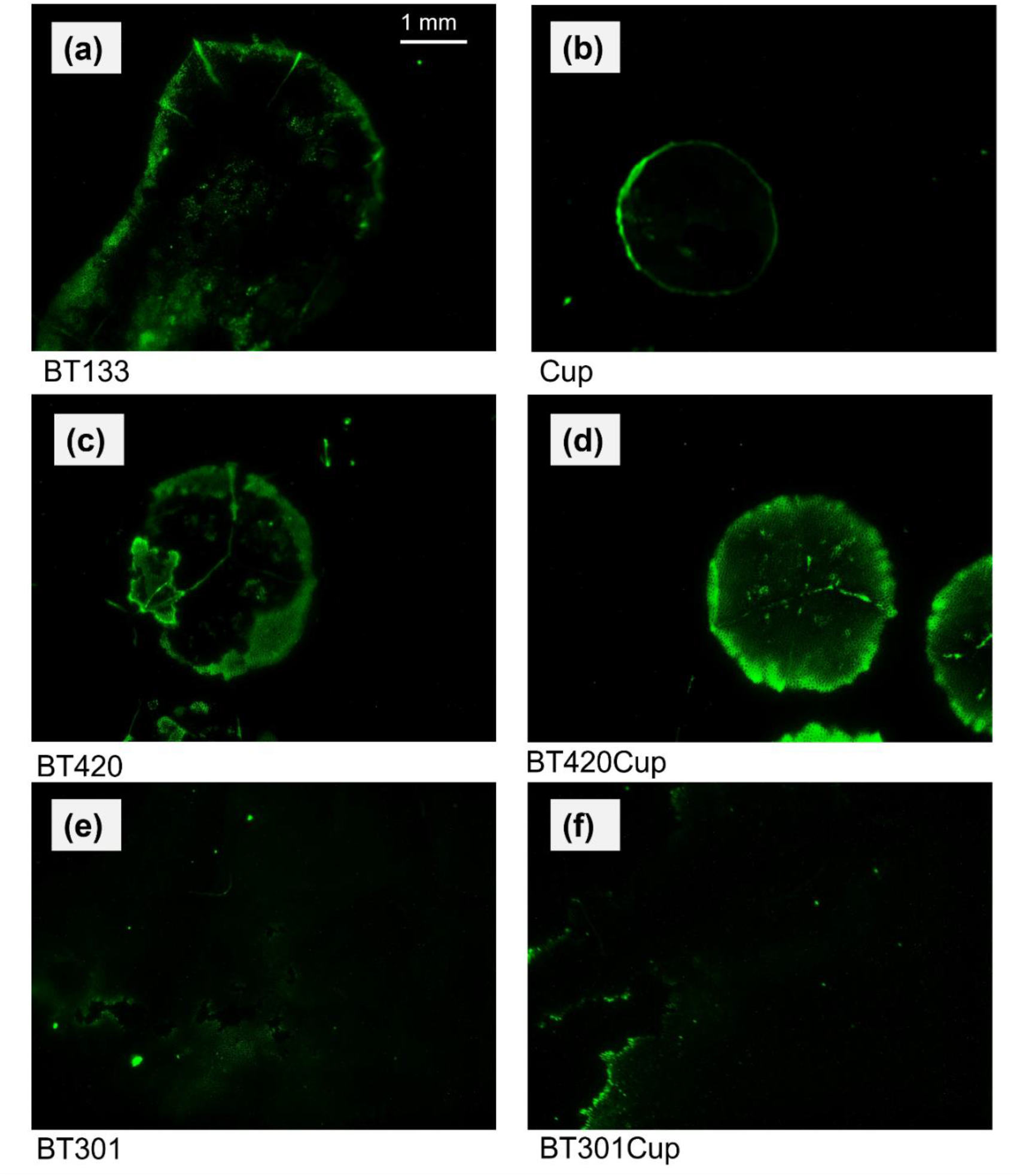
Distribution of copper formulation droplets with BREAK-THRU^®^ additives on the adaxial side of apple leaves. Droplet size was 5 μl. Leaves were misted with fluorescence marker (H2DCFDA) to trace the copper deposition and post-treatment leaf intra-cellular tissue response. Fluorescence pictures were recorded on treated surfaces using multispectral fluorescence microscopy. RGB images were taken at 20x magnification, covering an area of 36 mm^2^. Treatments were BT133 (a), Cup (b), BT420 (c), BT420Cup (d), BT301 (e), BT301Cup (f).

### 3.3 Phytotoxic potential and intra-cellular tissue activity

The electron transport rate (ETR, μmol m^-1^ s^-1^) was not affected by either treatment with copper alone or copper with BREAK-THRU^®^ additives at copper concentrations of 2 g L^-1^ and 4 g L^-1^, except for the Cup treatment alone on the adaxial side of apple leaves (Figure 3a, 3b, Supporting Information Table S5). At a higher copper concentration of 12 g L^-1^, ETR on the adaxial leaf side was significantly reduced for leaves treated with Cup alone or BT420Cup as compared to water-treated leaves (Supporting Information Table S5). The photochemical efficiency of photosystem II (Fv/Fm) was not negatively influenced by copper or additive treatment (Figure 3c, 3d, Supporting Information Table S6). The maximum fluorescence yield (Fm’) on the abaxial leaf side was reduced by BT301 and BT301Cup in several cases at concentrations of 2 g L^-1^ and 12 g L^-1^, but not at 4 g L^-1^ in the light-adapted state and by BT301 and BT301Cup at a concentration of 2 g L^-1^ in the dark-adapted state (Supporting Information Table S7). BT420Cup at copper concentrations of 2, 4, and 12 g L^-1^ and Cup and copper concentrations of 4 and 12 g L^-1^ reduced the quantum yield of nonregulated energy dissipation (NO) in the dark-adapted state on the adaxial leaf side (Supporting Information Table S8). NO in the light-adapted state and the quantum yield of regulated energy dissipation (NPQ) were not negatively influenced by treatments with BREAK-THRU^®^ tank-mixtures (Supporting Information Table S9).

**Figure 3:**
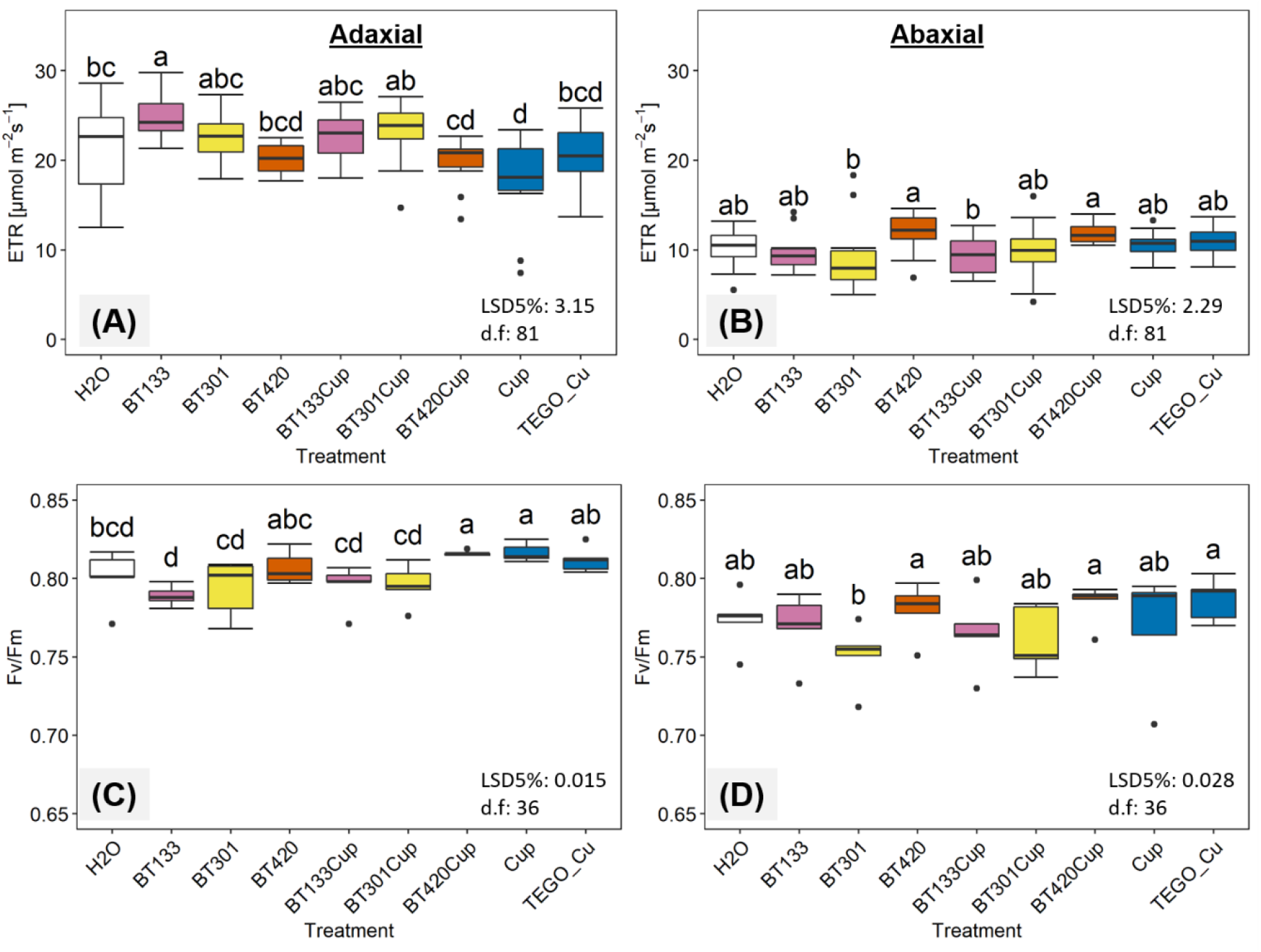
Electron transport rate (ETR, μmol m^-2^ s^-1^) in the light-adapted state (a, b) and photochemical efficiency of photosystem II in the dark-adapted state (Fv/Fm) (c, d) of apple leaves treated with copper formulations (dose 4 g L^-1^) with BREAK-THRU^®^ (BT133: 1.5 ml L^-1^, BT301: 0.75 ml L^-1^, BT420: 2.5 ml L^-1^) on adaxial and abaxial surfaces of apple leaves, respectively. ETR and Fv/Fm were measured on dark-adapted leaves with a chlorophyll fluorescence device (Imaging-PAM). Measurements was taken 7 days after treatment (abaxial) and 8 days after treatment (adaxial). Boxplots show median ± quartile (lower 25%, upper 75%). Different letters denote significant differences between ETR or Fv/Fm values (LSD 5%, least significant difference between two means at P = 0.05; d.f., degrees of freedom associated with LSDs, n = 5 in dark-adapted state, and n = 10 in light-adapted state).

Five days after treatment, leaves that were pre-treated with BT301 or BT301Cup and misted with H2DCFDA, showed an increase in relative fluorescence area, indicating higher ROS production as a signal of intra-cellular tissue activity. Other copper or BREAK-THRU^®^ treatments did not affect the fluorescence area (Figure 4).

**Figure 4:**
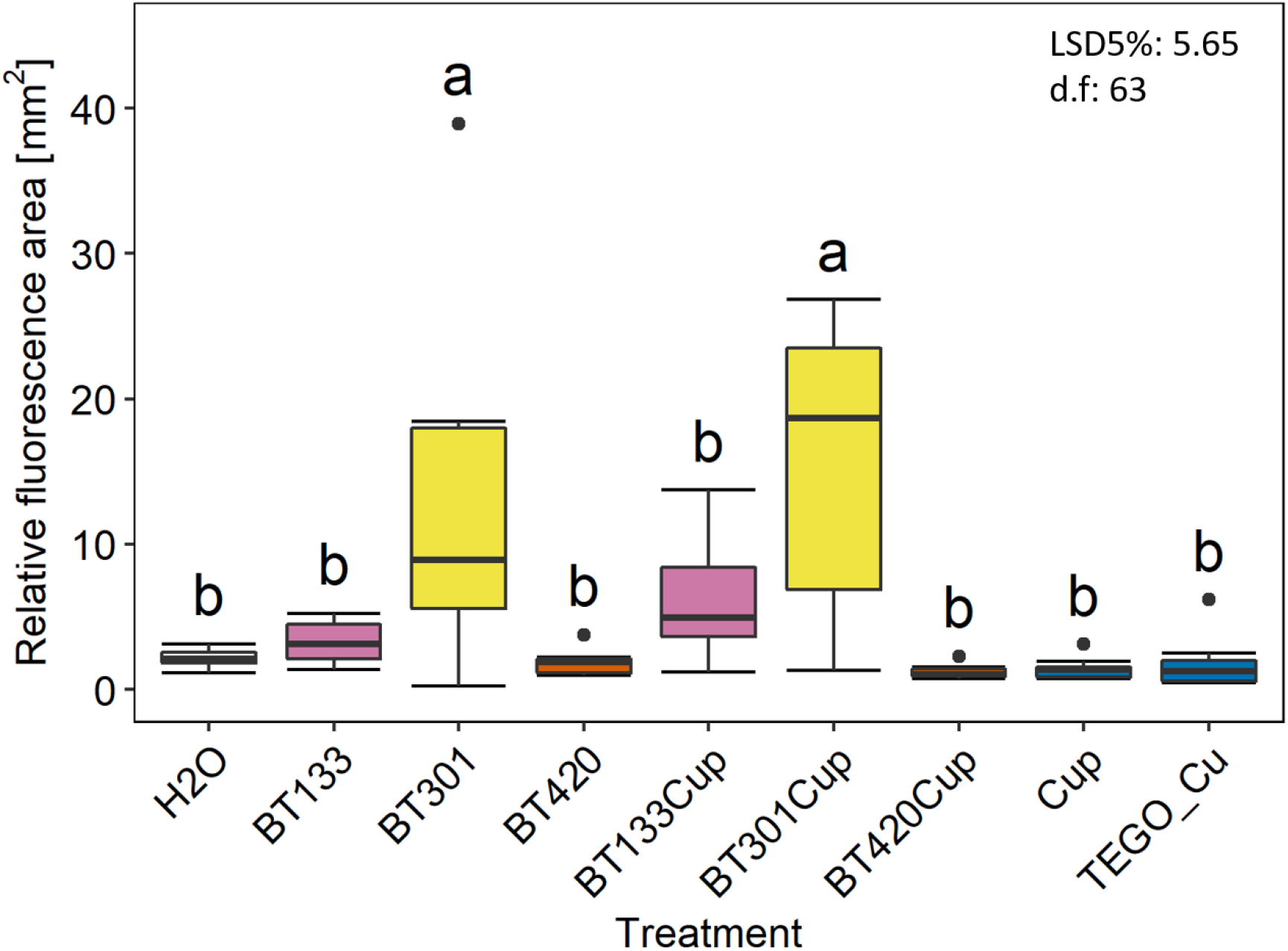
Relative fluorescence area (mm^2^) on apple leaves showing leaf intra-cellular tissue activity or ROS accumulation. The leaves were pre-treated with BREAK-THRU^®^ (BT133: 1.5 ml L^-1^, BT301: 0.75 ml L^-1^, BT420: 2.5 ml L^-1^) additives and copper preparations (1000 mg Cu^2+^ L^-1^) with BREAK-THRU^®^ additives. Leaves were misted with a fluorescence marker (H2DCFDA). Fluorescence images were recorded on treated surfaces using multispectral fluorescence microscopy with 11x magnification. The size of the measured leaf area was 124 mm^2^. Boxplots show median ± quartile (lower 25%, upper 75%). Different letters denote significant differences between relative fluorescence areas (LSD 5%, least significant difference between two means at P = 0.05; d.f., degrees of freedom associated with LSDs, n = 8).

### 3.4 Biological efficacy

Seven days after inoculation, 75% of the water-treated control leaves showed symptoms of infection by *V. inaequalis*, whereas leaves that were pre-treated with copper preparations (Cup and TEGO_Cu) or BT133Cup showed significantly lower infection intensity, and leaves treated with BT420Cup and BT301Cup were symptom-free (Figure 5a, Table 2). However, a few leaves treated with BT301Cup developed necroses, especially at leaf tips or leaf edges. Some leaves that were pre-treated with demineralised water, Cup or BT133Cup were additionally infected with *P. leucotricha* (Figure 5b, Table 2).

**Figure 5:**
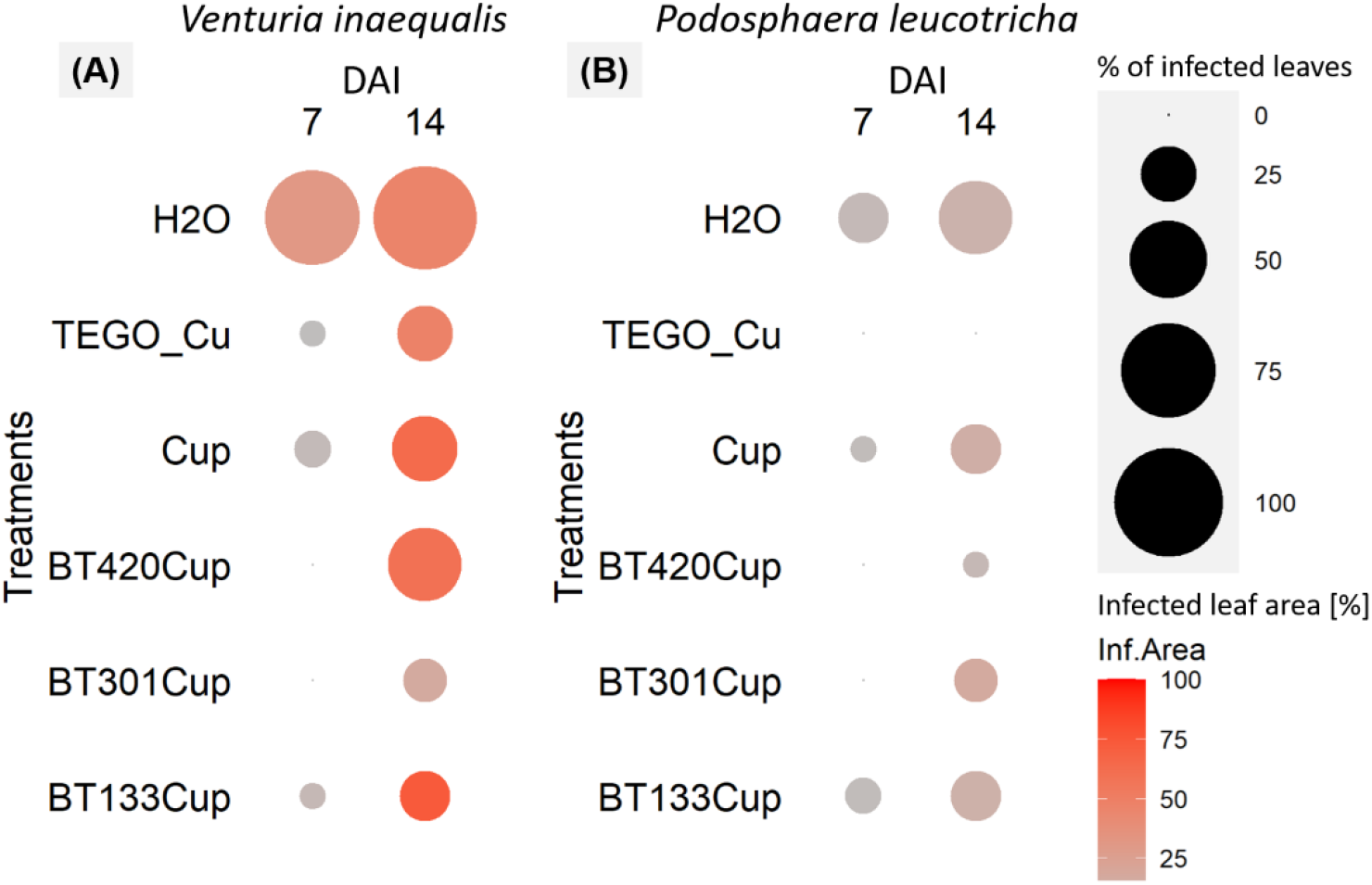
Percentage of infected apple leaves with visible *Venturia inaequalis* (a) or *Podosphaera leucotricha* (b) symptoms (circle size) and percentage of infected area on leaves (colour). Plants were pre-treated with copper and BREAK-THRU^®^ as tank mixtures. Assessments were done 7 and 14 days after inoculation (DAI). Sample size for each treatment was 20 (5 biologically replicated plants with 4 leaves per plant per treatment). Note: increasing diameter of a circle indicates increasing percentage of infected leaves. Increasing intensity of red signals signifies increasing percentage of infected leaf area.

**Table 2:** Percentage of infected leaves caused either by *Venturia inaequalis* or by *Podosphaera leucotricha*. Plants were pre-treated with copper and BREAK-THRU^®^ as tank mixtures. Visual symptom assessment was done 7 and 14 days after inoculation (DAI) on 5 biologically replicated plants with 4 leaves per plant per treatment (in total 20 leaves). Different letters denote significant differences between treatment groups (LSD 5%, least significant difference between two means at P = 0.05; d.f., degrees of freedom associated with LSDs; SEM = standard error of the mean).

| DAI | Treatment | Infected leaves [%] |  |  |  |
| --- | --- | --- | --- | --- | --- |
|  |  | *Venturia inaequalis* |  | *Podosphaera leucotricha* |  |
| | | (mean ± SEM) | | (mean ± SEM) | |
| 7 | H<sub>2</sub>O | 75 ± 19 | a | 20 ± 5 | a |
| 7 | Cup | 10 ± 6 | b | 10 ± 6 | ab |
| 7 | TEGO_Cu | 5 ± 5 | b | 0 ± 0 | b |
| 7 | BT133Cup | 0 ± 0 | b | 5 ± 5 | b |
| 7 | BT301Cup | 0 ± 0 | b | 0 ± 0 | b |
| 7 | BT420Cup | 0 ± 0 | b | 0 ± 0 | b |
| LSD5% |  | 24.9 |  | 11.1 |  |
| d.f |  | 24 |  | 24 |  |
| 14 | H<sub>2</sub>O | 90 ± 6 | a | 45 ± 9 | a |
| 14 | Cup | 40 ± 6 | bc | 20 ± 9 | ab |
| 14 | TEGO_Cu | 25 ± 8 | bcd | 0 ± 0 | b |
| 14 | BT133Cup | 20 ± 9 | cd | 20 ± 9 | ab |
| 14 | BT301Cup | 15 ± 10 | d | 15 ± 15 | b |
| 14 | BT420Cup | 45 ± 9 | b | 5 ± 5 | b |
| LSD5% |  | 24.2 |  | 27.0 |  |
| d.f |  | 24 |  | 24 |  |

Fourteen days after inoculation, 90% of the water-treated leaves showed symptoms of infection with *V. inaequalis*, whereas the shares of infected leaves were significantly lower for leaves pre-treated with Cup (35%) and TEGO_Cu (25%). Plants that were pre-treated with BT133Cup, BT420Cup or BT301Cup showed significantly lower infection intensity at 20%, 45% and 15% of leaves, respectively (Figure 5a, Table 2). Symptom spread on the infected leaves was strongest on leaves that were pre-treated with BT133Cup (74 ± 14%) and weakest on those pre-treated with BT301Cup (17 ± 3%) (Figure 5a). Infection with *P. leucotricha* was observed on 45% of untreated control leaves, 20% of leaves treated with BT133Cup or Cup, 15% of leaves treated with BT301Cup and 5% of BT420Cup-treated leaves. Leaves that were pre-treated with TEGO_Cu remained symptom-free (Figure 5b, Table 2).

## 4. Discussion

Reducing the internal surface tension of spray droplets is an important function of spreader adjuvants (Hazen, 2000). Adding BREAK-THRU^®^ additives to demineralised water led to a strong reduction in surface tension. For BT301, surface tension decreased from 72.14 ± 0.18 mN m^-1^ to 20.95 ± 0.18 mN m^-1^. BT420 also showed a considerable surface tension reduction to 37.83 ± 0.21 mN m^-1^, whereas the addition of BT133 had only a small or no effect on surface tension (Figure 1a, Supporting Information Table S1). This observation was supported by observations of contact angles, which responded similarly to the addition of additives to the adaxial and abaxial leaf side as well as to a smooth silicone surface (Figure 1b, Supporting Information Table S1, S2). Contact angles are not only influenced by the contacting fluids, but also by the structure and the composition of participating interfaces (Drelich et al., 2020). Contact angles of BT133 and BT420 were between 5.04° and 25.18° smaller on leaf surfaces than on the silicone surface, and contact angle measurement with BT301 was not possible at all on the apple leaf surface.

As tank-mix additives, BT301 and BT420 significantly reduced the contact angles of copper formulations (i.e., Funguran progress, Fung or Cuprozin progress, Cup) as compared to copper formulation alone (Figure 1f, Supporting Information Table S3). However, they did not show a further decreasing effect on contact angles when the copper concentration in the treatment increased (Figure 1f, Supporting Information Table S4). In contrast to the other two additives, the impact of BT133 on contact angles depended on the Cup concentration, with a reducing effect compared to Cup alone appearing on the adaxial leaf side at 4 and 12 g Cup L^-1^. This trend was associated with a decreasing surface tension (from 65.73 ± 0.38 mN m^-1^ at 2 g L^-1^ to 55.38 ± 0.20 mN m^-1^ at 12 g L^-1^) at increasing Cup concentrations (Figure 1e). The additives enhanced surface wetness and distribution of Cup treatments. Without additives, Cup droplets formed coffee-ring structures, with fragmented surface coverage (Figure 2b). Copper is accumulated in high concentrations within these rings. This may be problematic, since copper can be phytotoxic to plant leaves, if it accumulates on the plant surface (Lalancette & McFarland, 2007). The coffee-ring structures were associated with reduced ETR values and PS(II) in the light-adapted state on the adaxial leaf side of Cup-treated leaves (Figure 3a, Supporting Information Table S5, S6). Such copper depositions may prevent light from entering into the leaves and reaching the chloroplasts. This may reduce the efficiency of photosystem II as well as decreasing PS(II) light and ETR. BT420 enhanced the surface wetness of Cup by reducing the contact angles on the adaxial leaf side by about 29° at a concentration of 4 g L^-1^ in comparison to Cup without additive (Figure 1f). This was accompanied by a uniform Cup-distribution within the droplets (Figure 2d). BT301 reduced the surface tension of Cup 4 g L^-1^ by about 49 mN m^-1^ to 20.58 ± 0.26 mN m^-1^, enabling spreading on the leaf surface (Figure 1e, Figure 2f). These results indicate that different components in spray mixtures interact with each other. Reducing surface tension is a known mode of action of some additives, such as BREAK-THRU^®^ S233 and S240 (Basi et al., 2014).

Apple leaves that were treated with the additives without copper formulation showed no signs of phytotoxic potential based on plant photosynthetic activity, as expressed by ETR, Fv/Fm, NO and NPQ (Figure 3, Supporting Information Table S5, S6, S8, S9). We observed that the maximum fluorescence yield (Fm’) was lower for the BT301 and BT301Cup treatments (Supporting Information Table S7). However, the relative fluorescence area after misting with H2DCFDA, an indicator for ROS accumulation (Fichman et al., 2019), was relatively higher for BT301 (8.88 ± 2.77 mm^2^) and BT301Cup (15.59 ± 3.60 mm^2^) treatments as compared to the demineralised water treatment (2.15 ± 0.22 mm^2^) (Figure 4). In general, higher ROS accumulation is an indication of oxidative stress. On the other hand, ROS accumulation tends to regulate stress responses. Since a reduction in photosynthetic activity was not observed at the applied concentration of either BT301 or BT301Cup, this may be due to the non-toxic level of ROS accumulation. This result is supported by previous studies reporting positive effects of ROS in plants at low levels of oxidative stress (Huang et al., 2019; Mittler, 2017). In exceptional cases, BT301Cup triggered local necrosis at the tips or edges of some of the youngest leaves, where the treatment solution might have accumulated during spray application. Such an accumulation or dripping-off effect is sometimes unavoidable in practice due to the phyllotaxy of crop plants, where leaf surfaces may not be level, making the BT301Cup solution accumulate at the tip or edges of some young leaves. Spray application volumes should therefore be adjusted to the leaf area, to achieve full coverage but prevent droplet accumulation on the leaf tips or leaf margins.

Pre-treatment with copper-additive mixtures before inoculation with *V. inaequalis* significantly reduced post-inoculation disease symptoms. In terms of the biological efficacy against *V. inaequalis*, BT301 showed the best synergistic effect among all copper-additive mixtures, followed by BT133Cup and TEGO_Cu. On plants that were pre-treated with BT301Cup, the number of infected leaves (15% of the rated leaves) (Figure 5a, Table 2) and the area of visible symptoms on the infected leaves (17%) were comparatively low at 14 days after inoculation (Figure 5a). Interestingly, the younger leaves above the rated leaves of BT301Cup treated plants also remained healthier than the younger canopy of Cup-treated plants. BT301Cup showed extensive surface wetness (Figure 1e, Figure 2f). The addition of BT301 to the treatment ensured that the copper active ingredient covered the whole leaf surface. Therefore, the treatment might have contact to nearly all pathogen conidia, allowing the copper compounds to reduce the germination rate of these conidia (Montag et al., 2006). BT133Cup also showed high effectiveness against *V. inaequalis*, only permitting infection of about 20% of the rated leaves. However, the infected area on leaves treated with BT133Cup (74%) was significantly larger than for the BT301Cup treatment (17%). Potentially, the sticking effect of BT133 may have improved the distribution of copper ions on the leaf surface, allowing it to maximize coverage of the leaf area.

The apple powdery mildew pathogen *P. leucotricha* is transmitted via windborne spores and develops fastest at temperatures between 10 and 25 °C and at relative humidity >70% (Marine et al., 2010). In the growth chamber, temperature and relative humidity were often within this optimal range, with temperatures between 15.8 and 24.5 °C and a relative humidity up to 76% within 14 days after inoculation. Accordingly, *P. leucotricha* symptoms increased strongly between 7 and 14 days after inoculation (Figure 5b). Plants treated with water showed the strongest infection spread during the 14 days after inoculation, leading to 45% infected leaves. Plants treated with TEGO_Cu or BT420Cup were nearly free from visible powdery mildew symptoms at the 14^th^ day after inoculation. Although *P. leucotricha* is controlled with sulphur formulations, the use of the selected additives in this study as a tank-mixture with copper formulations aimed at controlling *V. inaequalis* might synergistically suppress disease symptoms caused by *P. leucotricha* (Table 2, Table 3). The synergistic effect of additives in tank-mix copper preparations may allow reducing the amount of copper applied in horticultural production by enhancing the biological efficacy of the copper preparation. Further studies should focus on detecting this synergistic effect under orchard conditions to evaluate the implications for horticultural practices in the field.

**Table 3:** Schematic summary of the selective performance of the tested additives in tank-mixtures based on substance properties, plant responses and biological activity (++ Strong positive effect, + positive effect, 0 neutral effect, - negative effect).

| Parameters \ Additives | BT133 | BT301 | BT420 |
| --- | --- | --- | --- |
| Surface wetness Cu | 0 | ++ | + |
| Synergetic effect with Cu against apple scab | + | ++ | + |
| Synergetic effect with Cu against powdery mildew | 0 | +/0 | ++ |
| Phytotoxicity | No | Possible at leaf tips or edges | No |

## 5. Conclusions

The results confirmed that tank-mix additives can reduce surface tension and contact angles on apple leaf surfaces, improving the distribution of Cu-preparations on plant surfaces. The selected additives and dosages appeared to cause no phytotoxic effects on apple leaves, yet efforts should be made to avoid the accumulation of BT301 droplets on leaf tips or edges. Our results imply that the additives may synergistically enhance the biological efficacy of Cu^2+^against plant pathogens (*V. inaequalis*, the causal agent of apple scab disease, and *P. leucotricha*, which causes apple powdery mildew) under controlled conditions. All three tank-mix additives (BT133, BT301 and BT420) have different synergistic effects with regard to surface wetness or in plant protection. Our observations indicated that BT301 may cause greater synergetic effects of Cu^2+^ against *V. inaequalis* than BT420, and BT420 may cause greater synergetic effects of Cu^2+^ against *P. leucotricha* than BT301.

## Abbreviations

BT133: BREAK-THRU^®^ SP 133
BT133Cup: Cuprozin^®^ progress with BREAK-THRU^®^ SP 133
BT133Fung: Funguran^®^ progress with BREAK-THRU^®^ SP 133
BT301: BREAK-THRU^®^ S 301
BT301Cup: Cuprozin^®^ progress with BREAK-THRU^®^ S 301
BT301Fung: Funguran^®^ progress with BREAK-THRU^®^ S 301
BT420: BREAK-THRU^®^ SF 420
BT420Cup: Cuprozin^®^ progress with BREAK-THRU^®^ SF 420
BT420Fung: Funguran^®^ progress with BREAK-THRU^®^ SF 420
CFI: Chlorophyll fluorescence imaging
Cup: Cuprozin^®^ progress
ETR: Electron transport rate (μmol m^-2^ s^-1^)
Fm’: Maximum fluorescence yield
Fung: Funguran^®^ progress
Fv/Fm: Photochemical efficiency of photosystem II
H2DCFDA: 2’,7’-dichlorofluorescin diacetate
NO: Quantum yield of nonregulated energy dispassion
NPQ: Quantum yield of regulated energy dispassion
PAR: Photoactive radiation (μmol m^-2^ s^-1^)
PS(II): Effective quantum yield of photosystem II
TEGO_Cu: TEGO^®^ XP 11052

## Author contributions

SP designed the experiments. CS and SP performed the experiments, collected data, analysed and prepared the manuscript. EL, CS, and SP reviewed, edited and agreed on the final manuscript.

## Acknowledgements

We thank Knut Wichterich, Getrudis Heimes and Harry Berg for their help on lab analysis and plant cultivation.

## Data availability statement

The data that support the findings of this study are available as supporting information.

## Ethical approval

This article does not contain any studies with human participants or animals performed by any of the authors.

