## Supporting Information_S1_to_S9 for "Synergism and phytotoxicity: the effects of tank-mix additives on the biological efficacy of Cu^2+^ against *Venturia inaequalis* and *Podosphaera leucotricha*"

**Supporting Information S1 – S9 (Schmitz et al)**

**Table_1_SuppInfo (S1):** Contact angles (°) and surface tension (mN $m^{-1}$) of different tank-mix additives dissolved in water on a silicone surface. Droplet size for contact angle was 5 µl, and left and right contact angles were merged. Different letters denote significant differences between treatment groups (LSD 5%, least significant difference between two means at P = 0.05; d.f., degrees of freedom associated with LSDs, SEM = standard error of the mean, n = 20 for contact angles, n = 10 for surface tension).

| **Treatment** | **Concentration [v/v]** | **Contact angle [°] (mean ± SEM)** | | **Surface tension [mN $m^{-1}$] (mean ± SEM)** | |
|---|---|---|---|---|---|
| $H_2O$ | 0 | 115.51 ± 0.28 | a | 72.14 ± 0.18 | a |
| BT133 | 0.015 | 111.62 ± 0.61 | b | 69.82 ± 0.40 | b |
| BT301 | 0.008 | 42.80 ± 0.58 | d | 20.95 ± 0.18 | d |
| BT420 | 0.025 | 96.09 ± 0.26 | c | 37.83 ± 0.21 | c |
| LSD 5% | | 1.30 | | 2.03 | |
| d.f | | 76 | | 36 | |

**Table_2_SuppInfo (S2):** Contact angles (°) of different tank-mix additives dissolved in water on the adaxial and abaxial leaf side of an apple leaf. Droplet size was 5 µl, and left and right contact angles were merged. Contact angle measurement of BT301 was not possible on apple leaves due to fast spreading. Different letters denote significant differences between treatment groups (LSD 5%, least significant difference between two means at P = 0.05; d.f., degrees of freedom associated with LSDs, SEM = standard error of the mean, n = 20).

| **Treatment** | **Concentration [v/v]** | **Adaxial (mean ± SEM)** | | **Abaxial (mean ± SEM)** | |
|---|---|---|---|---|---|
| $H_2O$ | 0 | 106.50 ± 0.86 | a | 101.32 ± 0.83 | b |
| BT420 | 0.025 | 70.91 ± 1.11 | b | 72.55 ± 0.78 | c |
| BT133 | 0.015 | 106.58 ± 0.41 | a | 104.19 ± 0.56 | a |
| LSD 5% | | 2.37 | | 2.08 | |
| d.f | | 57 | | 57 | |

**Table_3_SuppInfo (S3):** Contact angles (°) and surface tension (mN $m^{-1}$) of different copper preparations with BREAK-THRU® treatments dissolved in water on a silicone surface. Droplet size for contact angle was 5 µl, and left and right contact angles were merged. Different letters denote significant differences between treatment groups (LSD 5%, least significant difference between two means at P = 0.05; d.f., degrees of freedom associated with LSDs, SEM = standard error of the mean, n = 20 for contact angles, n = 10 for surface tension).

| **Treatment** | **Concentration [g $L^{-1}$]** | **Contact angle [°] (mean ± SEM)** | | **Surface tension [mN $m^{-1}$] (mean ± SEM)** | |
|---|---|---|---|---|---|
| $H_2O$ | 0 | 115.51 ± 0.28 | a | 72.14 ± 0.18 | a |
| Fung | 3 | 115.23 ± 0.49 | a | 70.40 ± 0.21 | b |
| TEGO_Cu | 4 | 112.2 ± 0.30 | b | 59.95 ± 0.46 | c |
| BT133Fung | 3 | 113.03 ± 0.38 | b | 60.33 ± 10.42 | c |
| BT301Fung | 3 | 61.43 ± 0.47 | d | 22.23 ± 0.15 | e |
| BT420Fung | 3 | 97.43 ± 0.58 | c | 41.60 ± 0.40 | d |
| LSD5% | | 1.21 | | 0.93 | |
| d.f | | 114 | | 54 | |

**Table_4_SuppInfo (S4):** Contact angles (°) of different copper preparations with BREAK-THRU® treatments dissolved in water on the adaxial and abaxial leaf side of an apple leaf. Droplet size was 5 µl, and left and right contact angles were merged. Contact angle measurement of BT301Cup was not possible on apple leaves due to fast spreading. Different letters denote significant differences between treatment groups, while performing multiple groups comparison (LSD 5%, least significant difference between two means at P = 0.05; d.f., degrees of freedom associated with LSDs, SEM = standard error of the mean, n = 20).

| Treatment | Concentration [g $L^{-1}$] | Adaxial (mean ± SEM) | | Abaxial (mean ± SEM) | |
|---|---|---|---|---|---|
| $H_2O$ | 0 | 106.50 ± 0.86 | a | 101.32 ± 0.83 | a |
| Cup | 2 | 99.58 ± 0.77 | c | 99.42 ± 0.80 | b |
| Cup | 4 | 100.76 ± 0.56 | c | 98.91 ± 0.56 | b |
| Cup | 12 | 100.31 ± 0.63 | c | 98.53 ± 0.36 | b |
| BT133Cup | 2 | 103.45 ± 0.38 | b | 101.42 ± 0.32 | a |
| BT133Cup | 4 | 93.03 ± 1.05 | e | 99.19 ± 0.63 | b |
| BT133Cup | 12 | 95.67 ± 0.49 | d | 91.55 ± 0.53 | c |
| BT420Cup | 2 | 73.01 ± 0.91 | g | 75.77 ± 0.75 | d |
| BT420Cup | 4 | 71.71 ± 0.93 | g | 70.13 ± 0.94 | e |
| BT420Cup | 12 | 77.09 ± 0.45 | f | 75.99 ± 0.33 | d |
| LSD5% | | 2.05 | | 1.79 | |
| d.f | | 190 | | 190 | |

**Table_5_SuppInfo (S5):** Electron transport rate (ETR, µmol $m^{-2}$ $s^{-1}$) in the light-adapted state of apple leaves treated with different doses (2 g $L^{-1}$, 4g $L^{-1}$ and 12 g $L^{-1}$) of copper formulations with BREAK-THRU® (BT133: 1.5 ml $L^{-1}$, BT301: 0.75 ml $L^{-1}$, BT420: 2.5 ml $L^{-1}$) on adaxial and abaxial surfaces of apple leaves, respectively. ETR was measured on dark-adapted leaves with a chlorophyll fluorescence device (Imaging-PAM). Measurement was done 7 days after treatment (abaxial) and 8 days after treatment (adaxial). (LSD 5%, least significant difference between two means at P = 0.05; d.f., degrees of freedom associated with LSDs, SEM = standard error of the mean, n= 10).

| **Treatments** | **Concentration [g $L^{-1}$]** | **ETR (µmol $m^{-2}$ $s^{-1}$)** Adaxial (mean ± SEM) | **Abaxial (mean ± SEM)** |
|---|---|---|---|
| $H_2O$ | 0 | 21.19 ± 1.59 | 10.521 ± 0.77 |
| BT133 | 0 | 24.76 ± 0.80 | 9.86± 0.72 |
| BT301 | 0 | 22.71 ± 0.90 | 9.40 ± 1.39 |
| BT420 | 0 | 20.14 ± 0.55 | 11.79 ± 0.76 |
| Cup | 2 | 17.58 ± 1.79 | 11.42 ± 0.47 |
| Cup | 4 | 17.39 ± 1.70 | 10.67 ± 0.47 |
| Cup | 12 | 17.23 ± 1.52 | 10.16 ± 0.84 |
| TEGO_Cu | 2 | 21.11 ± 0.91 | 12.06 ± 0.49 |
| TEGO_Cu | 4 | 20.45 ± 1.12 | 10.90 ± 0.51 |
| TEGO_Cu | 12 | 18.00 ± 1.07 | 11.25 ± 1.03 |
| BT133Cup | 2 | 24.27 ± 1.30 | 8.92 ± 0.81 |
| BT133Cup | 4 | 22.67 ± 0.81 | 9.29 ± 0.68 |
| BT133Cup | 12 | 22.81 ± 0.65 | 8.31 ± 0.68 |

| | | | |
|---|---|---|---|
| BT301Cup | 2 | 22.85 ± 1.35 | 10.56 ± 1.15 |
| BT301Cup | 4 | 22.91 ± 1.16 | 9.86 ± 1.11 |
| BT301Cup | 12 | 22.65 ± 1.07 | 10.08 ± 1.28 |
| BT420Cup | 2 | 20.39 ± 0.90 | 11.99 ± 0.43 |
| BT420Cup | 4 | 19.76 ± 0.93 | 11.84 ± 0.36 |
| BT420Cup | 12 | 17.49 ± 1.10 | 11.38 ± 0.55 |
| LSD5% | | 3.26 | 2.28 |
| d.f | | 171 | 171 |

**Table_6_SuppInfo (S6):** Photochemical efficiency of photosystem II in the dark-adapted (Fv/Fm) and light-adapted (PS(II)) state of apple leaves treated with different doses (2 g $L^{-1}$, 4g $L^{-1}$ and 12 g $L^{-1}$) of copper formulations with BREAK-THRU® (BT133: 1.5 ml $L^{-1}$, BT301: 0.75 ml $L^{-1}$, BT420: 2.5 ml $L^{-1}$) on adaxial and abaxial surfaces of apple leaves, respectively. Fv/Fm and PS(II) were measured on dark-adapted leaves with a chlorophyll fluorescence device (Imaging-PAM). Measurement was done 7 days after treatment (abaxial) and 8 days after treatment (adaxial). (LSD 5%, least significant difference between two means at P = 0.05; d.f., degrees of freedom associated with LSDs; SEM = standard error of the mean; n = 5 in dark-adapted state, and n = 10 in light-adapted state).

| Treatment | Conc. [g $L^{-1}$] | Fv/Fm (dark adapted state) Adaxial (mean ± SEM) | Fv/Fm (dark adapted state) Abaxial (mean ± SEM) | PS(II) (light adapted state) Adaxial (mean ± SEM) | PS(II) (light adapted state) Abaxial (mean ± SEM) |
|---|---|---|---|---|---|
| $H_2O$ | 0 | 0.800 ± 0.008 | 0.773 ± 0.008 | 0.210 ± 0.016 | 0.131 ± 0.010 |
| BT133 | 0 | 0.789 ± 0.003 | 0.769 ± 0.001 | 0.251 ± 0.010 | 0.128 ± 0.009 |
| BT301 | 0 | 0.794 ± 0.008 | 0.751 ± 0.009 | 0.228 ± 0.008 | 0.125 ± 0.018 |
| BT420 | 0 | 0.807 ± 0.005 | 0.780 ± 0.008 | 0.206 ± 0.006 | 0.151 ± 0.010 |
| Cup | 2 | 0.812 ± 0.002 | 0.781 ± 0.009 | 0.178 ± 0.018 | 0.144 ± 0.007 |
| Cup | 4 | 0.817 ± 0.003 | 0.769 ± 0.016 | 0.177 ± 0.017 | 0.136 ± 0.006 |
| Cup | 12 | 0.815 ± 0.002 | 0.778 ± 0.009 | 0.176 ± 0.015 | 0.131 ± 0.011 |
| TEGO_Cu | 2 | 0.808 ± 0.003 | 0.783 ± 0.006 | 0.215 ± 0.009 | 0.154 ± 0.005 |
| TEGO_Cu | 4 | 0.812 ± 0.004 | 0.787 ± 0.006 | 0.208 ± 0.011 | 0.141 ± 0.006 |
| TEGO_Cu | 12 | 0.809 ± 0.07 | 0.785 ± 0.007 | 0.185 ± 0.011 | 0.143 ± 0.011 |
| BT133Cup | 2 | 0.788 ± 0.004 | 0.762 ± 0.010 | 0.244 ± 0.013 | 0.119 ± 0.011 |
| BT133Cup | 4 | 0.795 ± 0.006 | 0.765 ± 0.011 | 0.233 ± 0.013 | 0.124 ± 0.009 |
| BT133Cup | 12 | 0.801 ± 0.004 | 0.766 ± 0.031 | 0.237 ± 0.006 | 0.114 ± 0.010 |
| BT301Cup | 2 | 0.792 ± 0.004 | 0.748 ± 0.007 | 0.229 ± 0.013 | 0.147 ± 0.018 |
| BT301Cup | 4 | 0.796 ± 0.006 | 0.761 ± 0.009 | 0.23 ± 0.012 | 0.129 ± 0.014 |
| BT301Cup | 12 | 0.799 ± 0.006 | 0.765 ± 0.005 | 0.226 ± 0.011 | 0.132 ± 0.017 |
| BT420Cup | 2 | 0.814 ± 0.002 | 0.777 ± 0.007 | 0.205 ± 0.010 | 0.155 ± 0.007 |
| BT420Cup | 4 | 0.816 ± 0.001 | 0.784 ± 0.006 | 0.206 ± 0.011 | 0.153 ± 0.005 |
| BT420Cup | 12 | 0.817 ± 0.001 | 0.783 ± 0.008 | 0.181 ± 0.011 | 0.146 ± 0.008 |
| LSD5% | | 0.013 | 0.026 | 0.033 | 0.030 |
| d.f | | 76 | 76 | 171 | 171 |

**Table_7_SuppInfo (S7):** Maximum fluorescence yield (Fm’) in the dark- and light-adapted state of apple leaves treated with different doses (2 g $L^{-1}$, 4 g $L^{-1}$ and 12 g $L^{-1}$) of copper formulations with BREAK-THRU® (BT133: 1.5 ml $L^{-1}$, BT301: 0.75 ml $L^{-1}$, BT420: 2.5 ml $L^{-1}$) on adaxial and abaxial surfaces of apple leaves, respectively. Fm’ was measured on dark-adapted leaves with a chlorophyll fluorescence device (Imaging-PAM). Measurement was done 7 days after treatment (abaxial) and 8 days after treatment (adaxial). (LSD 5%, least significant difference between two means at P = 0.05; d.f., degrees of freedom associated with LSDs; SEM = standard error of the mean; n = 5 in dark-adapted state, and n = 10 in light-adapted state).

| Treatment | Conc. [g $L^{-1}$] | Fm‘ (dark adapted state) | | Fm‘ (light adapted state) | |
|---|---|---|---|---|---|
| | | Adaxial (mean ± SEM) | Abaxial (mean ± SEM) | Adaxial (mean ± SEM) | Abaxial (mean ± SEM) |
| $H_2O$ | 0 | 0.368 ± 0.018 | 0.464 ± 0.019 | 0.107 ± 0.004 | 0.226 ± 0.012 |
| BT133 | 0 | 0.370 ± 0.009 | 0.414 ± 0.028 | 0.124 ± 0.005 | 0.229 ± 0.012 |
| BT301 | 0 | 0.390 ± 0.039 | 0.327 ±10.023 | 0.137 ± 0.015 | 0.158 ± 0.009 |
| BT420 | 0 | 0.388 ± 0.019 | 0.483 ± 0.030 | 0.136 ± 0.012 | 0.235 ± 0.007 |
| Cup | 2 | 0.412 ± 0.025 | 0.456 ± 0.018 | 0.143 ± 0.011 | 0.237 ± 0.016 |
| Cup | 4 | 0.406 ± 0.012 | 0.467 ± 0.023 | 0.147 ± 0.011 | 0.247 ± 0.017 |
| Cup | 12 | 0.399 ± 0.020 | 0.446 ± 0.017 | 0.130 ± 0.009 | 0.215 ± 0.011 |
| TEGO_Cu | 2 | 0.393 ± 0.015 | 0.464 ± 0.019 | 0.155 ± 0.073 | 0.268 ± 0.019 |
| TEGO_Cu | 4 | 0.403 ± 0.015 | 0.467 ± 0.005 | 0.156 ± 0.019 | 0.257 ± 0.026 |
| TEGO_Cu | 12 | 0.365 ± 0.014 | 0.413 ± 0.016 | 0.137 ± 0.023 | 0.215 ± 0.017 |
| BT133Cup | 2 | 0.338 ± 0.017 | 0.418 ± 0.023 | 0.108 ± 0.005 | 0.210 ± 0.017 |
| BT133Cup | 4 | 0.367 ± 0.017 | 0.416 ± 0.015 | 0.118 ± 0.006 | 0.216 ± 0.017 |
| BT133Cup | 12 | 0.345 ± 0.018 | 0.369 ± 0.038 | 0.112 ± 0.006 | 0.188 ± 0.013 |
| BT301Cup | 2 | 0.345 ± 0.012 | 0.298 ± 0.026 | 0.114 ± 0.004 | 0.149 ± 0.015 |
| BT301Cup | 4 | 0.352 ± 0.011 | 0.393 ± 0.042 | 0.116 ± 0.003 | 0.180 ± 0.018 |
| BT301Cup | 12 | 0.338 ± 0.011 | 0.376 ± 0.045 | 0.114 ± 0.006 | 0.194 ± 0.030 |
| BT420Cup | 2 | 0.387 ± 0.011 | 0.451 ± 0.020 | 0.133 ± 0.011 | 0.223 ± 0.019 |
| BT420Cup | 4 | 0.37 ± 0.006 | 0.459 ± 0.027 | 0.131 ± 0.008 | 0.222 ± 0.020 |
| BT420Cup | 12 | 0.379 ± 0.017 | 0.412 ± 0.037 | 0.131 ± 0.010 | 0.208 ± 0.022 |
| LSD5% | | 0.050 | 0.075 | 0.033 | 0.049 |
| d.f | | 76 | 76 | 171 | 171 |

**Table_8_SuppInfo (S8):** Quantum yield of nonregulated energy dissipation (NO) in dark- and light-adapted state of apple leaves treated with different doses (2 g $L^{-1}$, 4g $L^{-1}$ and 12 g $L^{-1}$) of copper formulations with BREAK-THRU® (BT133: 1.5 ml $L^{-1}$, BT301: 0.75 ml $L^{-1}$, BT420: 2.5 ml $L^{-1}$) on adaxial and abaxial surfaces of apple leaves, respectively. NO was measured on dark-adapted leaves with a chlorophyll fluorescence device (Imaging-PAM). Measurement was done 7 days after treatment (abaxial) and 8 days after treatment (adaxial). (LSD 5%, least significant difference between two means at P = 0.05; d.f., degrees of freedom associated with LSDs; SEM = standard error of the mean; n = 5 in dark-adapted state, and n = 10 in light-adapted state).

| **Treatment** | **Conc. [g $L^{-1}$]** | **NO (dark adapted state)** | | **NO (light adapted state)** | |
|---|---|---|---|---|---|
| | | **Adaxial (mean ± SE)** | **Abaxial (mean ± SE)** | **Adaxial (mean ± SE)** | **Abaxial (mean ± SE)** |
| $H_2O$ | 0 | 0.200 ± 0.008 | 0.227 ± 0.008 | 0.232 ± 0.012 | 0.422 ± 0.020 |
| BT133 | 0 | 0.211 ± 0.003 | 0.231 ± 0.010 | 0.252 ± 0.008 | 0.484 ± 0.022 |
| BT301 | 0 | 0.206 ± 0.008 | 0.249 ± 0.009 | 0.269 ± 0.014 | 0.434 ± 0.025 |
| BT420 | 0 | 0.193 ± 0.005 | 0.220 ± 0.008 | 0.275 ± 0.014 | 0.415 ± 0.008 |
| Cup | 2 | 0.188 ± 0.002 | 0.219 ± 0.009 | 0.283 ± 0.016 | 0.449 ± 0.038 |
| Cup | 4 | 0.183 ± 0.003 | 0.231 ± 0.016 | 0.296 ± 0.021 | 0.462 ± 0.039 |
| Cup | 12 | 0.185 ± 0.002 | 0.222 ± 0.009 | 0.268 ± 0.016 | 0.426 ± 0.034 |
| TEGO_Cu | 2 | 0.192 ± 0.003 | 0.217 ± 0.006 | 0.303 ± 0.041 | 0.498 ± 0.044 |
| TEGO_Cu | 4 | 0.188 ± 0.004 | 0.213 ± 0.006 | 0.303 ± 0.034 | 0.473 ± 0.048 |
| TEGO_Cu | 12 | 0.191 ± 0.007 | 0.215 ± 0.007 | 0.297 ± 0.040 | 0.452 ± 0.042 |
| BT133Cup | 2 | 0.212 ± 0.004 | 0.238 ± 0.010 | 0.244 ± 0.013 | 0.441 ± 0.029 |
| BT133Cup | 4 | 0.205 ± 0.006 | 0.235 ± 0.011 | 0.247 ± 0.012 | 0.451 ± 0.030 |
| BT133Cup | 12 | 0.199 ± 0.003 | 0.234 ± 0.013 | 0.248 ± 0.011 | 0.457 ± 0.028 |
| BT301Cup | 2 | 0.208 ± 0.004 | 0.252 ± 0.007 | 0.255 ± 0.007 | 0.422 ± 0.029 |
| BT301Cup | 4 | 0.204 ± 0.006 | 0.239 ± 0.009 | 0.255 ± 0.008 | 0.396 ± 0.024 |
| BT301Cup | 12 | 0.201 ± 0.006 | 0.235 ± 0.005 | 0.260 ± 0.012 | 0.436 ± 0.040 |
| BT420Cup | 2 | 0.186 ± 0.002 | 0.223 ± 0.007 | 0.271 ± 0.019 | 0.416 ± 0.031 |
| BT420Cup | 4 | 0.184 ± 0.001 | 0.216 ± 0.006 | 0.280 ± 0.016 | 0.403 ± 0.028 |
| BT420Cup | 12 | 0.183 ± 0.001 | 0.217 ± 0.008 | 0.278 ± 0.013 | 0.428 ± 0.032 |
| LSD5% | | 0.013 | 0.026 | 0.056 | 0.090 |
| d.f | | 76 | 76 | 171 | 171 |

**Table_9_SuppInfo (S9):** Quantum yield of regulated energy dissipation (NPQ) in the light-adapted state of apple leaves treated with different doses (2 g $L^{-1}$, 4g $L^{-1}$ and 12 g $L^{-1}$) of copper formulations with BREAK-THRU® (BT133: 1.5 ml $L^{-1}$, BT301: 0.75 ml $L^{-1}$, BT420: 2.5 ml $L^{-1}$) on adaxial and abaxial surfaces of apple leaves, respectively. NPQ was measured on dark-adapted leaves with a chlorophyll fluorescence device (Imaging-PAM). Measurement was done 7 days after treatment (abaxial) and 8 days after treatment (adaxial). (LSD 5%, least significant difference between two means at P = 0.05; d.f., degrees of freedom associated with LSDs; SEM = standard error of the mean; n = 10).

| **Treatment** | **Concentration [g $L^{-1}$]** | **NPQ** | |
|---|---|---|---|
| | | **Adaxial (mean ± SE)** | **Abaxial (mean ± SE)** |
| $H_2O$ | 0 | 0.519 ± 0.017 | 0.434 ± 0.011 |
| BT133 | 0 | 0.497 ± 0.009 | 0.388 ± 0.020 |
| BT301 | 0 | 0.503 ± 0.011 | 0.442 ± 0.016 |
| BT420 | 0 | 0.519 ± 0.017 | 0.434 ± 0.011 |
| Cup | 2 | 0.539 ± 0.021 | 0.407 ± 0.037 |
| Cup | 4 | 0.527 ± 0.016 | 0.402 ± 0.036 |
| Cup | 12 | 0.556 ± 0.021 | 0.443 ± 0.033 |
| TEGO_Cu | 2 | 0.482 ± 0.043 | 0.349 ± 0.044 |
| TEGO_Cu | 4 | 0.490 ± 0.030 | 0.387 ± 0.049 |
| TEGO_Cu | 12 | 0.518 ± 0.039 | 0.405 ± 0.046 |
| BT133Cup | 2 | 0.513 ± 0.024 | 0.441 ± 0.021 |
| BT133Cup | 4 | 0.520 ± 0.010 | 0.425 ± 0.028 |
| BT133Cup | 12 | 0.516 ± 0.016 | 0.429 ± 0.035 |
| BT301Cup | 2 | 0.516 ± 0.010 | 0.431 ± 0.013 |
| BT301Cup | 4 | 0.515 ± 0.009 | 0.475 ± 0.013 |
| BT301Cup | 12 | 0.514 ± 0.008 | 0.432 ± 0.025 |
| BT420Cup | 2 | 0.524 ± 0.027 | 0.429 ± 0.030 |
| BT420Cup | 4 | 0.513 ± 0.023 | 0.444 ± 0.025 |
| BT420Cup | 12 | 0.541 ± 0.020 | 0.427 ± 0.027 |
| LSD5% | | 0.061 | 0.084 |
| d.f | | 171 | 171 |
